# Circular RNAs exhibit limited evidence for translation, or translation regulation of the mRNA-counterpart in terminal hematopoiesis

**DOI:** 10.1101/2020.10.22.348276

**Authors:** Benoit P. Nicolet, Sjoert B.G. Jansen, Esther Heideveld, Willem H. Ouwehand, Emile van den Akker, Marieke von Lindern, Monika C. Wolkers

## Abstract

Each day, about 10^12^ erythrocytes and platelets are released into the blood stream. This substantial output from hematopoietic stem cells is tightly regulated by transcriptional and epigenetic factors. Whether and how circular RNAs (circRNAs) contribute to the differentiation and/or identity of hematopoietic cells is to date not known. We recently reported that erythrocytes and platelets contain the highest levels and numbers of circRNAs amongst hematopoietic cells. Here, we provide the first detailed analysis of circRNA expression during erythroid and megakaryoid differentiation. CircRNA expression not only significantly increased upon enucleation, but also had limited overlap between progenitor cells and mature cells, suggesting that circRNA expression stems from regulated processes rather than resulting from mere accumulation. To study circRNA function in hematopoiesis, we first compared the expression levels of circRNAs with the translation efficiency of their mRNA-counterpart. We found that only 1 out of 2531 (0.04%) circRNAs associated with mRNA-translation regulation. Furthermore, irrespective of 1000s of identified putative open reading frames, deep ribosome-footprinting sequencing and mass spectrometry analysis provided little evidence for translation of endogenously expressed circRNAs. In conclusion, circRNAs alter their expression profile during terminal hematopoietic differentiation, yet their contribution to regulate cellular processes remains enigmatic.

## INTRODUCTION

Each hour, millions of red blood cells (RBC) and platelets are produced in the bone marrow and released into the blood stream. Their production is mediated from erythroid and megakaryoid progenitors. The generation of these progenitors from hematopoietic stem cells is a highly orchestrated process. Tight regulation is obtained by the coordinated expression of transcription factors, long non-coding RNAs and micro-RNAs (1–3). Recently, we showed that also circular RNAs (circRNAs) are abundantly expressed in hematopoietic cells, and that their expression alters during differentiation (4).

CircRNAs are single stranded, circular RNA molecules that are generated from pre-mRNA transcripts through back-splicing (5–7). Back-splicing is driven by the classical spliceosome machinery at canonical splice sites (8), and circularization is mediated by pairing of complementary sequences, typically Alu elements, which are found in flanking introns. The pairing of Alu elements is regulated by RNA-binding proteins such as ADAR, DHX9, and NF90 (9–11). CircRNA can contain both intron and exon segments. While intronic circRNA are mostly retained in the nucleus (12), exonic circRNA undergo nuclear translocation in a size dependent manner (13, 14). In the cytoplasm, circRNAs are found to be exonuclease-resistant (15). In addition, circRNA lack the 3’ poly-adenylation tail, which renders them more stable than linear RNA (16).

Many functions have been attributed to circRNAs. CircRNAs were suggested to act as transcriptional activators (12, 17), or to serve as RNA-binding protein cargo (18). CircRNAs can also act as miRNA ‘sponge’, a feature that is to date limited to about ten circRNAs genome-wide in humans (19–21). In addition, circRNA-mediated regulation of mRNA translation has been proposed in cell lines (22–24). Lastly, it was shown with over-expression constructs in cell lines (25, 26), and from exemplary circRNAs in drosophila and human cell lines that circRNA can be translated into protein (27, 28). How these findings translate to endogenous circRNA expression during human hematopoietic differentiation, is not known.

We and others have observed that RBCs and platelets had the highest numbers and highest circRNA content of all analyzed mature cell types in humans (4, 29, 30). However, the origin of these high circRNA levels is not known, and has thus far been attributed to transcriptome degradation (30). In addition, the function of these circRNA in blood cell development remains unknown.

In the study presented here, we provide a detailed and comprehensive analysis of circRNA expression in erythroid and megakaryoid development. We identified >20,000 circRNAs in erythroid differentiation, and >12,000 in megakaryoid differentiation. The expression of circRNAs alters dramatically upon enucleation and platelet formation, and only partially (~40%) follows mRNA expression in differentiating blood cells. Integration of RNA-seq and ribo-seq data revealed that circRNA-mediated regulation of mRNA translation is not the main mode of action during megakaryocytic maturation. Furthermore, ribosome-footprinting analysis and mass spectrometry analysis demonstrated that translation of endogenous circRNAs may occur in reticulocytes, megakaryocytes, and/or platelets, however this is not a frequent event. In conclusion, even though circRNAs are highly prevalent in erythroid and megakaryoid development, their mode of action and activity in hematopoietic cells does not follow the proposed models and thus remains to be defined.

## MATERIALS AND METHODS

### CircRNA identification and analysis

Data on erythroblast differentiation were retrieved from our previous study (31). circRNA identification was performed with previously established circRNA detection pipeline (4). Briefly, the quality of RNA-seq reads was assessed using FastQC version 0.11.8 (Simon Andrews, Babraham institute). Data were aligned to the human genome (GRCh37 / hg19-release75) using STAR version 2.5.2b (32) allowing for chimeric detection, with an overhang ‘anchor’ of 15 nt on either side of the read. The chimeric output file of STAR was analyzed with DCC 0.4.6 (33) and CircExplorer2 (CE2) 2.3.2 (34) to detect, filter, and annotate circRNA. DCC was used for detection of linear reads at the circRNA coordinates (using the option -G). CircRNA expression was considered low confidence when at least 2 junction reads were found in at least 1 sample by both DCC and CE, and high confidence detection when at least 2 junction reads were found in all biological replicates of one specific cell type, by both tools. Supporting circRNA read counts were normalized to reads per million mapped reads (RPM). The maximal circRNA spliced length was calculated with the exon length information provided by the CE2 annotations. These annotations were then used to calculate the first and last circularized exon. For differential expression analysis of high confidence circRNA, we used DESeq2 (35) with p-adjusted <0.05. We used the total number of mapped reads per sample to calculate the scaling factors, instead of the automatic scaling factors detection of DESeq2. The log2 Fold Change (LFC) was calculated using the formula: *LFC(AB) = log2(B) - log2(A)*. Data analysis was performed in R (3.5.1) and R-Studio (1.1.453). Gene ontology of circRNA-expressing gene was performed using pantherDB.

### RNA-seq analysis of mRNA and ncRNA

After quality control with FastQC, raw RNA-seq reads were aligned with Salmon version 0.13.1 (36) on the coding and non-coding transcriptome (ENSEMBL, GRCh38 release 92) to obtain Transcripts Per kilobase per Million (TPM) normalized counts. To get mRNA and ncRNA expression per gene, TPM counts were summed up per gene name, according to ENSEMBL BioMart annotations (37).

### In-vitro megakaryocyte differentiation

Megakaryocytes were cultured from human cord blood-derived CD34+ cells. Cord blood was diluted in PBS, loaded on ficoll-pague (GE Healthcare) and centrifuged 15 minutes at 400g. The white blood cell layer was collected from the ficoll gradient, washed twice in PBS+ 1mM EDTA + 0.2% HSA. CD34+ cells were purified with anti-CD34 Microbeads (Miltenyi biotec 130-046-702) and MACS LS column (Miltenyi, 130-042-401) according to manufacturer’s guidelines. Purity for CD34+ cells was >90%, as determined by flow-cytometry with anti-CD34 antibody (#581-PE; Beckman Coulter, measured on Beckman Coulter FC500). CD34+ cells were seeded at a density of 50.000 cells/well in a 12 well plate in CellgroSGM (CellGenix) supplemented with 100ng/mL human recombinant TPO (CellGenix, Cat.1417) and 10ng/mL IL1β (Miltenyi Biotec, Cat.130-093-895). At day 3, 1:1 Fresh media was added with an additional 10ng/mL TPO and 1ng/mL IL1β. On day 5, cells were split at a density of 100.000 cells in 6 wells plates, and fresh media with 10ng/mL TPO and 1ng/mL IL1β was added. On day 10, cells were harvested and supplemented with 100μg/mL Cycloheximide (Sigma, Cat. 1810-1G) in culture medium prior to cell sorting.

### Megakaryocyte selection

Cultures cells were harvested when >85% of cells were CD41a+ (CD41a-APC; clone HIP8; BD Pharmingen; Cat.559777), as determined by flow-cytometry. Cells were pelleted at 120g for 1 minute at 4°C and washed once in ice cold buffer (PBS, 1mM EDTA, 0.2% HSA + 100μg/mL cycloheximide). Two-step selection was performed on ice with EasySep® Human PE-Positive Selection Kit (18551), using anti-human CD42b-PE (clone HIP1, BD Pharmingen, 555473) according to manufacturer’s guidelines. CD41a+ CD42b- and CD41a+ CD42b+ cell fractions were selected, washed using PBS + 100μg/mL cycloheximide, and kept on ice until further processing. Viability of the cells was >90% (7-AAD, Biolegend), and purity for CD41a+ CD42b- was >85% and CD41a+ CD42b+ was >95%, as determined by flow-cytometry. Cells were immediately processed.

### RNA-sequencing

Cell lysis and RNA isolation for RNA sequencing and ribosomal footprinting was performed according to Ignolia et al. (38) with some adaptations. Briefly, CD42- and CD42+ selected megakaryocytes were pelleted. Cell pellets were dissolved in 150μL ice cold lysis buffer (20mM Tris pH 7.4, 250mM NaCl2, 5mM MgCl2, 0.5% Triton X-100, 1mM Dithiotreitol (DTT), 0.024U/μL Turbo DNAse (Ambion), 0.01U/μL RNAseOUT (Invitrogen) and 100μg/mL Cycloheximide). The lysate was immediately split in two aliquots of 50 and 100 μL, for total RNA-sequencing and ribosomal footprint analysis, respectively. 2.5μL SuperaseIN RNAse inhibitor (Invitrogen) was immediately added to the total RNA sample, and total RNA was extracted using the miRNAeasy kit (Qiagen) according to manufacturer’s protocol. RNA quality was assessed on Agilent bioanalyzer RNA chip (all samples with RIN >9.0), and quantified using Qubit 2.0 (Thermofisher). Ribosomal RNA was removed using Ribo Zero Gold (Epicentre/Illumina; 20020596) using the manufacturer’s protocol, and the cDNA library was constructed with the KAPA stranded RNA-seq kit (KAPA biosystems KK8400) according to manufacturer’s protocol with Illumina Truseq forked adapters. Library quality was assessed on Agilent Bioanalyzer high sensitivity DNA chip and quantified using qPCR with the KAPA library quantification kit (KK4824). Libraries were pooled to a 2nM concentration and 125bp paired-end reads were sequenced on HiSeq 2000 (Illumina).

### Ribosomal footprinting

100 μL of cell-lysates was used for ribosome foot-printing preparation. Samples were kept on ice for 10 minutes and regularly vortexed to fully lyse the outer membranes and release the ribosomes. Lysates were spun at 20.000g for 10min at 4°C and supernatant was harvested to remove the remaining pelleted cell debris and nuclei. Lysates were treated with 2μL of RNAse 1 (Ambion) and incubated at 37°C for 45 minutes. The reaction was stopped by the addition of 5μL SUPERAseIN RNAse inhibitor. Lysates were loaded on a 1M 0.150mL Sucrose cushion in 8×34 mm polycarbonate thick wall tubes and centrifuged at 120.000g for 30 minutes in a TLA-120.1 rotor in a pre-cooled 4°C table top ultracentrifuge. The pellet was resuspended using 100μL resuspension buffer (10mM Tris pH 7.0, 1% SDS and 0.01U/μL proteinase K (NEB)) for 30 minutes at 37°C. Footprints were subsequently extracted with miRNAeasy kit according to manufacturer’s guidelines for small RNA. rRNA contamination was removed using the Ribo-Zero gold rRNA removal kit (epicenter/Illumina; 20020596). Footprint RNA quality was assessed using the Agilent Bioanalyzer using the RNA chip. Footprints were selected for 26-34nt with custom markers (see (38)) on Elchrom Scientific ORIGINS system and short fragment gels, using gel electro-elution into Spectra/Por 3 3.5KD MWCO dialysis membranes. Libraries were produced using the Art-Seq kit (Epicentre/Illumina) according to the manufacturer’s protocol. The quality of libraries was assessed on Agilent Bioanalyzer with high sensitivity DNA chip and quantified using qPCR with the KAPA library quantification kit (KK4824). Libraries were pooled to a 2nM concentration and sequenced on HiSeq 2000 (Illumina).

### Ribo-seq analysis

The quality of raw ribo-seq reads was assessed using FastQC. Reads were trimmed for sequencing adapters using Trimmomatic version 0.39 (39) or cutadapt version 3.0 (40), excluding reads <25 and >35nt long. Additional quality control was performed on the BAM files resulting from genome mapping with STAR in Ribo-seQC (41).

Trimmed reads were mapped with Salmon on the human transcriptome (GRCh38 release 92) to get TPM normalized counts. Using a k-mer of 11 performed best for ribo-seq mapping to build the Salmon index. Translation efficiency was obtained by dividing the normalized ribo-seq counts in TPM by the normalized RNA-seq counts in TPM. To estimate the ribosome footprint read density, we used the linearized sequence of high confidence circRNA and the coding transcriptome. After aligning the ribo-seq reads with Salmon, ribo-seq reads density per kilobase were obtained by dividing the estimated counts (not TPM) by circRNA length (for circRNA) and CDS length (for mRNA). The circular-over-linear ribo-seq reads density per kilobase (ribo-CLR) was obtained by dividing the circRNA ribo-seq density per kilobase, by the ribo-seq density per kilobase of mRNA. The ratio was calculated separately for each sample and averaged per population. Ribo-seq data for erythroblast and platelets were retrieved from a previous study (42).

### CircRNA-specific detection from ribo-seq reads

To identify circRNA-specific ribo-seq reads, we used an approach similar to the ‘classical’ circRNA detection described above. Genome alignment as described above with STAR was used for chimeric reads detection with incremental anchors of 8 to 16 nt on either side of a chimeric read, allowing for 1 mismatch. CE2 was then used to detect, quantify and annotate the circRNAs, and data were further processed in R.

### ORF prediction

The linearized, spliced ‘high confidence’ circRNA sequences were obtained using BEDtools version 2.18 (43). The sequence was juxtaposed 3 time to mimic a circRNA sequence. Open reading frames (ORFs) were predicted for both linearized circRNA sequence (1xLinRNA) and tripled (3xCircRNA) sequences using ORF-finder (version 0.4.3; NCBI). Resulting ORF sequences were analyzed and filtered in R.

### Detection of circRNA peptides by mass spectrometry

The high confidence circRNA sequences including all exons between first and last exon were obtained using BEDtools. The last 77nt of the circRNA sequences were joined to the first 77nt to generate a junction sequence library. This library was translated in 3 frames. Open reading frames (ORFs) that presented a length <30 amino acids (that is, not spanning the circRNA junction) were removed. All ORFs were transformed into a fasta library. A proteome reference library containing circRNA junction ORFs, as well as circRNA-specific ORFs (from 3xCircRNA sequences of platelets, filtered for unique circ-specific ORFs), and Uniprot Human proteome reference (downloaded 13-02-2019), was constructed and used for peptide matching with Proteome Discoverer (version 2.2; Thermo-Fisher) of our previously published mass spectrometry dataset on platelets from healthy donor (44). The precursor mass tolerance was set to 10ppm with a fragment mass tolerance of 0.6Da and a target FDR of 0.01. Only peptides with high confidence detection in MaxQuant were considered.

#### Plots and graphs

Heatmaps were generated in R using corrplot (version 0.84; T. Wei and V. Simko, 2017, https://github.com/taiyun/corrplot) or pheatmap 1.0.8 (45). Plots and graphs were generated with ggplot2 (46) in R, or Graphpad PRISM version 7.0.

### Data retrieval and deposition

For *ex vivo* RBC and platelet analysis, we used our previously published data sets. Data were retrieved from National Center for Biotechnology Information (NCBI) Gene Expression Omnibus (GEO) and Sequence Repository Archive (SRA) accession numbers for platelets (project: PRJEB4522): ERR335311, ERR335312 and ERR335313 (47); for RBCs: (GEO: GSE63703) SRR2124299, SRR2124300, SRR2124301 and (GEO: GSE69192) SRR2038798 (30, 48) ; for erythroid differentiation: GEO: GSE124363 (31); and for human cell lines GEO: GSE125218 (49).

Ribosome footprinting datasets of platelets and erythroblasts were retrieved from GEO with accession number: GSE85864 (42), and for those of human cell lines GEO: GSE125218 (49). For comparison to previous study of our transcriptome data, we retrieved additional datasets from HSC and MEP (50); cultured megakaryocytes (51); pro-, early/late basophilic, poly- and ortho-chromatic erythroid cells (52).

Raw mass spectrometry datasets of platelets were obtained from the PRoteomics IDEntifications Database (PRIDE) repository under the accession number PXD009020 (44).

## RESULTS

### CircRNA are abundant during erythropoiesis

We first investigated if and how circRNA expression alters during terminal erythropoiesis. To this end, we examined paired-end RNA sequencing data of erythroblasts isolated from 4 donors that were cultured for 12 days under differentiating conditions, and that were harvested on each day between day 0 to 7, in addition to day 9 and 12 (Figure 1A;(31)). This *in vitro* differentiation model resulted in overall gene expression patterns that closely resembled that of *ex vivo* isolated cells ((Figure S1A; Table S1; (31)). Also physical properties such as hemoglobin content, oxygen association and dissociation capacity and deformability matched those of *ex vivo* red blood cells (31).

**Figure 1:**
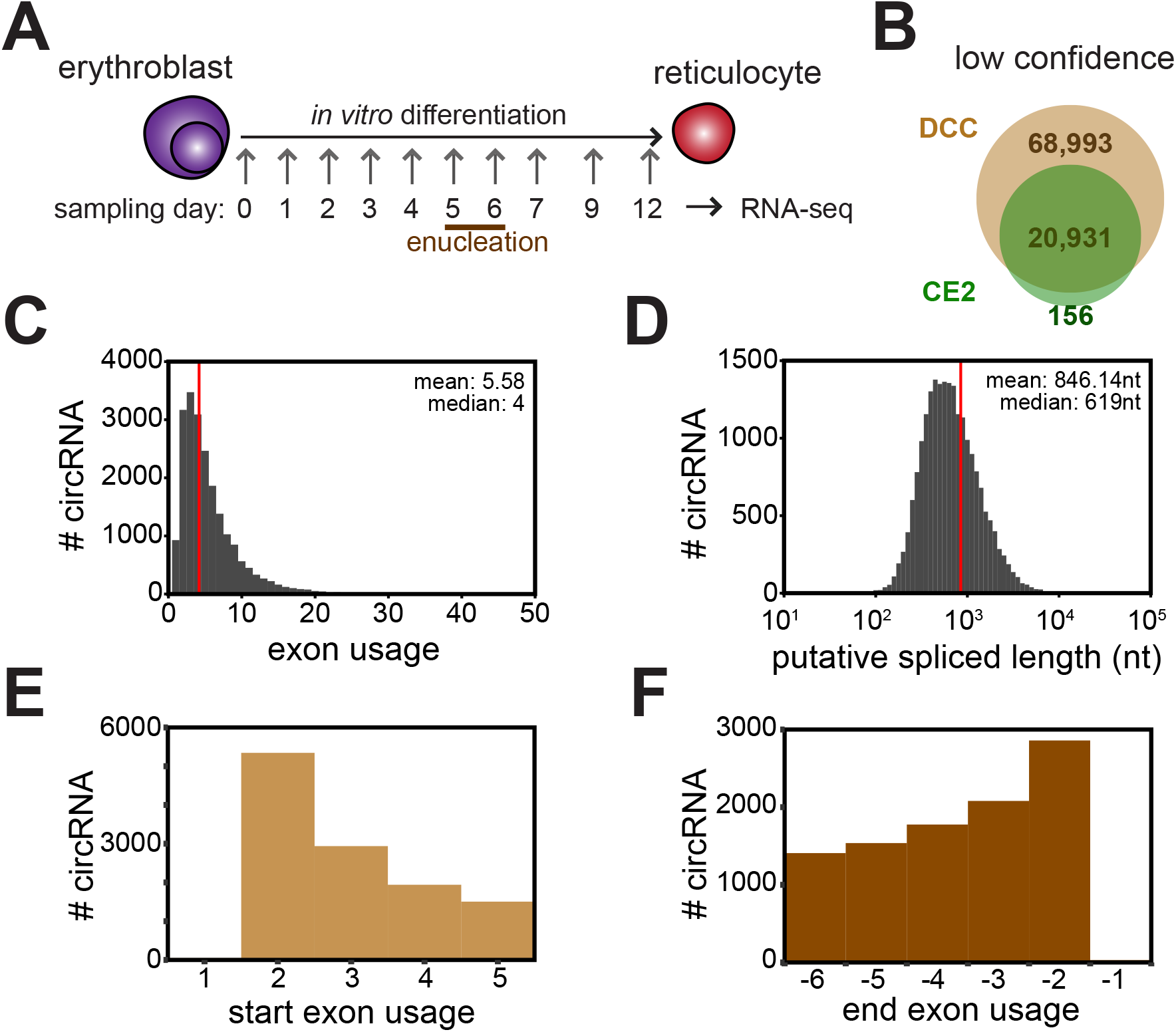
CircRNAs are abundant during *in vitro* erythroid differentiation. (**A**) Erythroblast progenitors were cultured for 12 days under differentiation conditions as in (31). Samples were taken at indicated days of differentiation, and RNA-seq was performed (n=4 donors). (**B**) CircRNA detection using DCC and CircExplorer2 (see methods) to identify ‘low confidence’ (circRNA with at least 2 reads in at least 1 sample) in all compiled samples from (A). The intersection of the circles represents the co-detected circRNAs with both tools. (**C**-**D**) Characterization of low confidence circRNA. (**C**) Putative exon usage and (**D**) putative spliced length, using splicing annotations from linear isoforms. (**E-F**) Characterization of (**E**) start and (**F**) end exon usage by ‘low confidence’ circRNAs, based on the circRNA junction positions and canonical mRNA splicing annotation.

The average sequencing depth of 30 million reads identified on average 27.3 million mapped reads. To detect, quantify, and to annotate circRNAs from chimeric reads, we used our established pipeline (4) that is based on the two algorithms DCC (33) and CircExplorer2 CE2, ((34); see methods section)). With a ‘low-confidence’ cut-off of at least 2 reads in at least one sample at any time of harvest, 20,931 circRNAs were co-detected by DCC and CE2 (Figure 1B; Table S2). Based on mRNA splicing annotations and circRNA junction reads, the estimated number of exons/circRNA was 5.58 in differentiating erythroblasts (median= 4 exons/circRNA; Figure 1C). The putative length of circRNAs was 846.14 nt (median= 619 nt; Figure 1D). In line with our previous work (4), circRNAs preferentially use the second exon of a linear transcript as the first circular exon (Figure 1E), a bias that is not as pronounced for the last circularized exon (Figure 1F). Thus, circRNAs are abundantly expressed during erythropoiesis.

### CircRNA levels increase at enucleation

To define how circRNA expression related to the expression of other transcripts during erythroid differentiation, we quantified the overall expression of protein-coding transcripts (mRNA), non-coding transcripts (ncRNA) and circRNA transcripts (TPM>0.1). We detected 19,449 mRNA-, 10,753 ncRNA-, and 4,526 circRNA-producing genes (Figure 2A). We also quantified the diversity of transcripts expressed during erythrocyte differentiation. At the early time points (i.e. up do day 5), erythroblasts mainly expressed mRNA and ncRNA transcripts (Figure 2B). These numbers steadily decreased over time and were reduced by 82.96% and 76.45%, respectively, at day 12 compared to day 0 (Figure 2B). In contrast, the number of circRNA transcripts substantially increased by a ~3.8-fold from day 6 onwards compared to day 0, and circRNA transcripts even surpassed the number of ncRNA transcripts at day 8 (Figure 2B). This steep increase of circRNAs coincided with the enucleation rate (Figure 2C).

**Figure 2:**
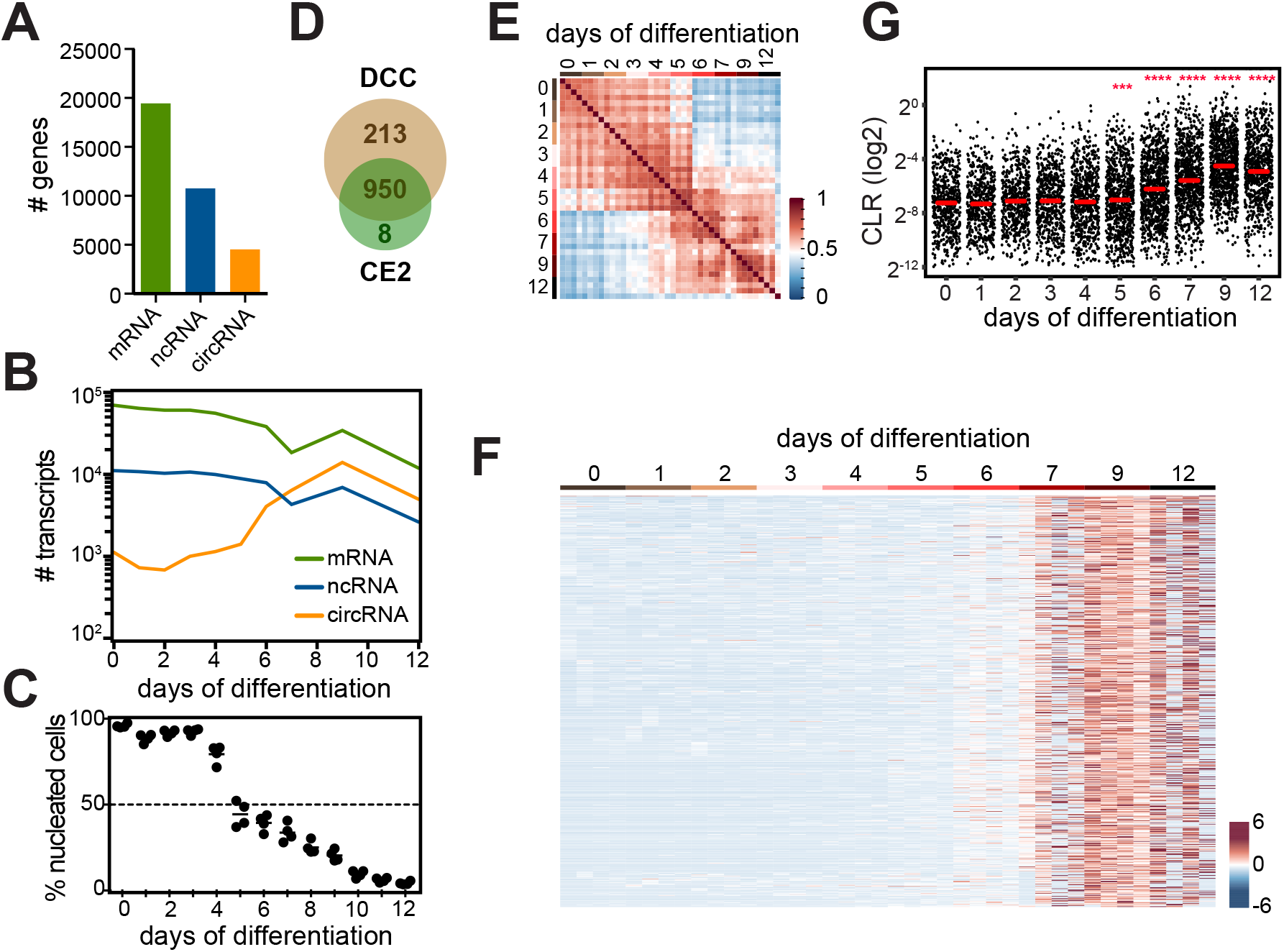
CircRNA expression increases at enucleation of erythroid cells. (**A**) Number of genes encoding mRNA, non-coding RNA (ncRNA) and circRNA, and (**B**) the number of transcripts detected during erythroid differentiation (cut-off: >0.1 TPM for mRNA and ncRNA; low confidence circRNA). (**C**) Percentage of enucleated cells at indicated time point (n=4 donors; adapted from (31)). (**D**) CircRNA detection using DCC and CircExplorer2 (see methods) from RNA-seq data generated from Figure 1. The intersection of the circle represents the co-detected circRNA referred to as ‘high confidence’ (circRNA with at least 2 reads in all 4 biological replicates of 1 time-point). (**E**) Pearson’s sample-correlation coefficient map of ‘high confidence’ circRNA expression. (**F**) Heatmap of the differentially expressed ‘high confidence’ circRNA (n=708; p-adjusted <0.05) corrected for sequencing depth (RPM). RPM: Reads per million mapped (linear) reads. (**G**) Circular over linear ratio (CLR) was calculated for ‘high confidence’ circRNAs during erythropoiesis, median is indicated in red. Differences were assessed by two-sided t test with Benjamini-Hochberg p-value adjustment, all compared to day 0. (***: padj<0.001, ****: padj<0.0001).

To measure the differential expression of circRNAs during erythropoiesis, we focused on ‘high confidence’ circRNAs, that is, a given circRNA is detected at least twice in all 4 donors at an individual time-point. This filter identified 950 circRNAs that were co-detected by DCC (n=1163) and CE2 (n=958; Figure 2D; Table S2). Gene ontology analysis of high confidence circRNAs revealed functions related to metabolic processes, catalytic and signaling activity (Figure S1B). Pearson’s correlation coefficient of all measured time points during erythropoiesis revealed 2 main clusters of circRNAs that were primarily expressed early (day 0-5), or late during differentiation (day 6-12; Figure 2E). In fact, a heatmap representation of high confidence circRNA expression showed a massive increase in circRNA expression from day 6 onwards (Figure S1C). When we compared all days to day 0 (Figure 2F), 708 circRNAs (74.5%) were differentially expressed (p-adjusted<0.05; Table S2). Of these, all but 1 circRNA, *circ-FIRRE* (Figure S1D), significantly increased their expression levels from day 6 onwards (Figure 2F; Table S2).

We next investigated how circRNA expression related to the expression of their mRNA counterparts in erythroid differentiation. To this end, we quantified the mRNA read counts at the circRNA start and end genomic position, using the build-in linear detection of DCC, as described previously (4). The circular-over-linear expression ratio (CLR) was calculated for each ‘high confidence’ circRNA, averaged across the 4 biological replicates for each time-point (Figure S1E; Table S2). Between day 0 and day 5 of erythropoiesis, the mRNA expression was higher compared to circRNA, as no circRNAs showed a CLR above 1. Only from day 6 onwards, 25 circRNAs showed a CLR>1, that is, circRNA expression is higher than that of their linear counterpart. Also the overall CLR increased from day 6 onwards and peaked at day 9 with a 7.3-fold increase compared to day 0 (Figure 2G). In conclusion, circRNA expression substantially alters at enucleation and can reach levels that are higher than those of their mRNA counterpart.

### CircRNA expression alters from megakaryocytes to platelets

We next investigated how circRNAs are expressed during megakaryopoiesis and in platelets, which are formed and released into the bloodstream by mature megakaryocytes. We performed RNA-seq analysis of three donors on purified immature CD41a^+^ CD42b^−^ (CD42-) and mature CD41a^+^ CD42b^+^ (CD42+) megakaryocytes that were cultured from cord blood CD34^+^ progenitors (see methods; Figure 3A). Again, also the gene expression of cultured MKs closely resembled that of previous studies (Figure S1A; Table S1). The average sequencing depth of 84.4 million reads per sample resulted in on average 74.6 million mapped reads to the genome. CircRNA detection with DCC and CE2 co-detected 12,809 circRNAs with low-confidence filtering (Figure 3B; Table S3). To define the relation of mRNA, ncRNA and circRNA expression during MK differentiation, we quantified the number of genes expressing these transcripts. We also included our previous analysis of circRNA expression in platelets to this analysis pipeline (4). Similar to erythropoiesis, the number of expressed mRNA and ncRNA genes decreased in particular in anucleate platelets, where the number of circRNA expressing genes increased by a 2.1-fold compared to MKs (Figure S2A). Furthermore, with 47,654 low confidence circRNAs detected, platelets had the highest diversity of circRNA transcripts (Figure 3C). In megakaryocytes, the estimated exon usage with a mean of 4.57 exon/circRNA (median= 4) and the putative spliced length with a mean of 663.9 nt/circRNA (median= 483nt), was slightly lower than that of circRNAs in erythropoiesis (Figure 1C, D, S2B, C). Yet, a preference for the 2^nd^ exon usage for circularization was apparent (29.2% of circRNA), with no overt preference of the last exon (14.2% of circRNA; Figure S2D, E).

**Figure 3:**
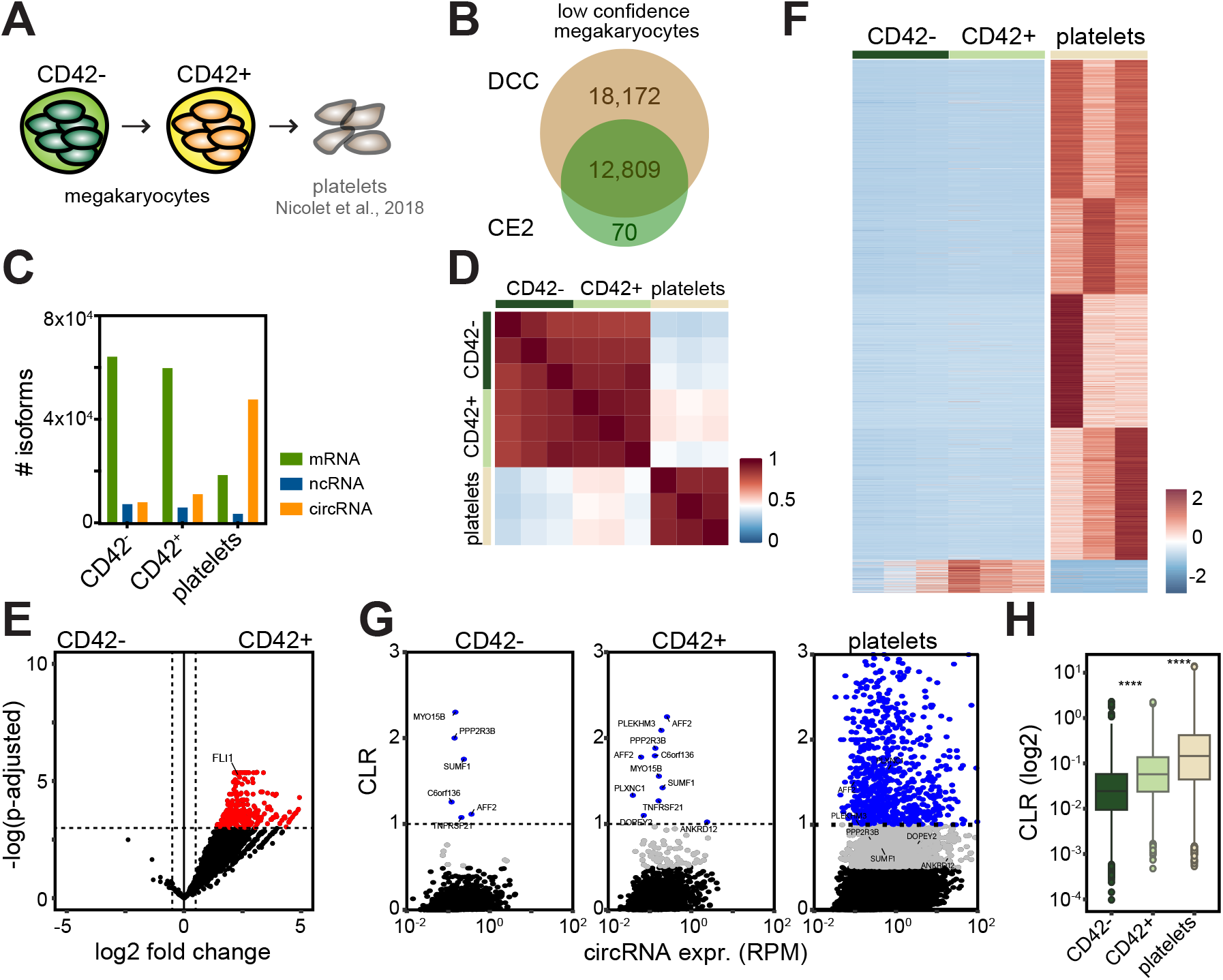
CircRNA expression during megakaryocyte differentiation. (**A**) Diagram representing *in vitro* differentiation from CD42- and CD42+ megakaryocytes (n=3 per population) to platelets (4). (**B**) CircRNA detection in CD42- and CD42+ megakaryocytes from RNA-seq data, using DCC and CircExplorer2. The intersection of the circle represents the co-detected ‘low confidence’ circRNAs. Of note, platelet data were not included for circRNA detection in megakaryocytes. (**C**) Number of mRNA, non-coding RNA (ncRNA) and circRNA transcripts detected during megakaryoid differentiation (>0.1 TPM for mRNA and ncRNA; low confidence circRNA). (**D**) Pearson’s sample-correlation coefficient map between CD42- and CD42+ megakaryocytes, and platelets (high confidence circRNA expression). (**E**) Volcano plot showing the differential expression of high confidence circRNA between CD42- (left) and CD42+ megakaryocytes (right; n=287; p-adjusted<0.05 and LFC>0.5, Z-score of reads per million mapped reads). (**F**) Heatmap of high confidence circRNAs detected in megakaryocytes and platelets. (**G**-**H**) Circular-over-linear ratio (CLR) was calculated for CD42- and CD42+ megakaryocytes and plotted against circRNA expression (**G**) or (**H**) compared between populations. Differences were assessed in H by two-sided t test p-value adjustment using the Benjamini-Hochberg procedure; ****: padj<0.0001.

To determine the differential circRNA expression in MKs, we focused on the 2531 high-confidence circRNAs (Figure S2F). Similar to erythroid cells, gene ontology analaysis of high confidence circRNAs in MKs showed a relation to metabolic processes, catalytic and signaling activity (Figure S2G). Pearson’s sample-correlation coefficient revealed a strong kinship between CD42- and CD42+ MK (Figure 3D). Yet, differential expression analysis of circRNA, identified 287 out of 2531 (11.34%) circRNAs, including the key transcription factor for MK differentiation FLI1 (p-adjusted<0.05 and log2 fold change >0.5; Figure 3E; Table S3). All differentially expressed circRNA were found in CD42+ MK. Platelets, however, showed an overall distinct expression pattern of circRNAs compared to the two megakaryocytic populations (Figure 3D, F; Table S3). Interestingly, not only the overall expression of circRNAs increased as CD42- MK matured towards platelets, but the overall CLR increased by ~2.1-fold during MK differentiation and by 3.11 between CD42+ MK and platelets ((4); Figure 3G, H; Table S3). Thus, circRNA are expressed and increase as megakaryocytes mature, yet substantially differ from circRNA expression in platelets.

### CircRNA expression can be independent of mRNA expression

We next studied how changes in circRNA expression related to changes in expression of the linear mRNA counterpart. To this end, we calculated the log2 fold change (LFC) in circRNA and mRNA expression in erythroid cells. We isolated five groups based on circRNA and mRNA expression changes (LFC>0.5): *group 1*: decreased in circRNA levels and decreased in mRNA levels; *group 2*: increased circRNA levels and decreased mRNA levels; *group 3*: increased circRNA levels and unchanged mRNA levels; *group 4*: increased circRNA levels and increased mRNA levels; *group 5*: unchanged circRNA levels; *group 6*: decreased circRNA and increased mRNA levels. When we compared the circRNA expression with the mRNA expression before enucleation, i.e. day 0 with day 5, we found that of 37.6% circRNAs, the changes (up or down) coincided with that of the mRNA counterpart (Figure 4A; group 1 (down) and 4 (up)). Of the 35.4% circRNAs that increased their expression (group 2 and 3), 18.3% of the mRNA counterpart decreased its expression (Figure 4A; group 2), and 17.1% remained unchanged (group 3). We also detected circRNAs whose expression levels remained unchanged throughout the first 5 days of differentiation (group 5; 24.6%).

**Figure 4:**
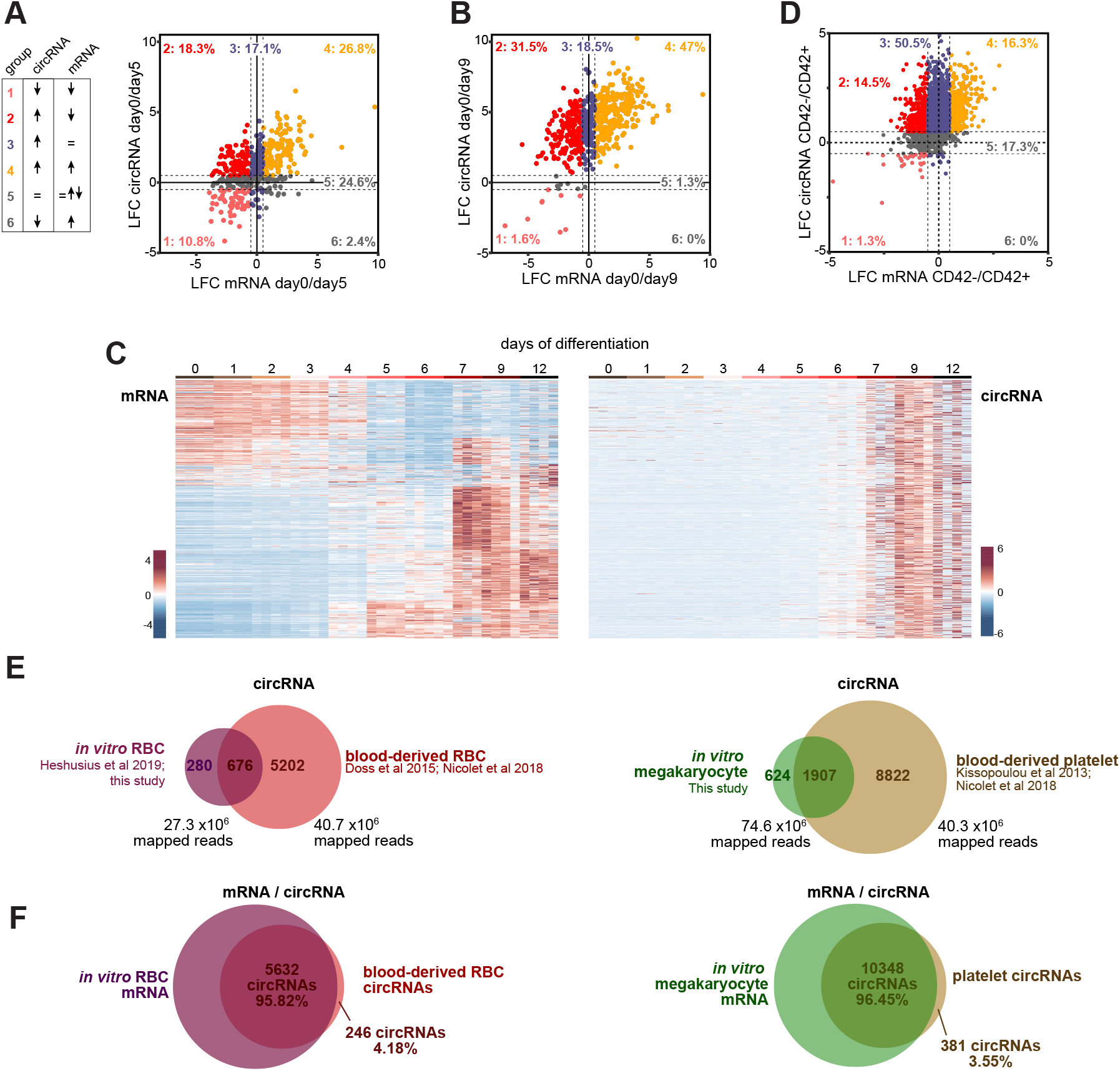
CircRNA expression can be independent of mRNA expression. (**A-B**) Log2 Fold Change (LFC) for circRNA and corresponding mRNA was calculated for (**A**) day 0 to 5 and (**B**) day 0 to 9 of erythroid differentiation, CircRNA-mRNA pairs are represented in: decreased circRNA / decreased mRNA (group 1, pink), increased circRNA / decreased mRNA (group 2, red), increased circRNA / equal mRNA (group 3, purple), increased circRNA / increased mRNA (group 4, orange), equal circRNA (group 5, grey), decreased circRNA / increased mRNA (group 6, grey). (**C**) Heatmap of mRNA expression and matched circRNA expression for all high confidence circRNA (n=950, Z-score of reads per million mapped reads). Note that mRNA expression was detected using DCC at the circRNA position (see methods). (**D**) LFC in circRNA and corresponding mRNA expression was calculated CD42- to CD42+ megakaryocytes differentiation. (**E**) Overlap in circRNA expression between *in vitro* RBC and *ex vivo* RBC (data from (4, 31, 48); left panel; Figure 1), and *in vitro* megakaryocytes and *ex vivo* platelets ((4, 47); right panel). Average sequencing depth is indicated under each population. (**F**) Overlap between circRNA in differentiated cells and mRNA in progenitor cells, for erythroid (left panel) and megakaryoid cells (right panel).

We then asked how the circRNA and mRNA expression relates upon enucleation. We compared the LFC of circRNA and mRNA expression of day 0 with that of day 9 of erythrocyte differentiation (Figure 4B). The overall high confidence circRNA expression increased at day 9 compared to day 0 by ~20 times from 32 to 651 identified circRNAs (Figure 4B). Owing to this increase, only 1.3% of circRNA did not alter, and 1.6% decreased their expression levels (Figure 4B; group 5 and 1, respectively). 48.6% of circRNAs showed an increase or decrease of expression in line with their mRNA counterpart (Figure 4B; group 1 and 4). Close to half (50.1%) of the circRNAs increased their expression levels, whereas the mRNA expression of the linear counterpart decreased or remained unchanged (Figure 4B; group 2; 31.5%, and group 3; 18.5%, respectively). When comparing day 5 or day 9 to day 0 of differentiation, we detected only 13 and 0 circRNAs, respectively, with decreased circRNA expression levels for which the mRNA levels increased (Figure 4A-B; group 6). We next questioned how the changes in circRNA expression related to mRNA changes measured at the circRNA positions throughout the entire erythroid differentiation process. We used the mRNA expression to define clusters, and matched these to the circRNA expression (Figure 4C). Whereas mRNA expression is more heterogeneously expressed, circRNA expression primarily increased around enucleation around day 6 (Figure 4C). Thus, the majority of circRNA do not follow the expression pattern of their linear counterpart in differentiating RBCs prior and after enucleation, indicating that circRNA expression in differentiating erythroid cells can be independent of mRNA expression.

We also compared the changes in circRNA and mRNA expression between CD42- and CD42+ megakaryocytes. The magnitude of the overall expression changes in MK was lower compared to erythroblasts (~7.6 LFC mRNA and ~7.7 LFC circRNA in MK, compared to ~16.7 LFC mRNA and ~13.9 LFC circRNA in erythroblasts from day 0 to 9; Figure 4A-D). 17.3% of circRNA did not alter in expression between CD42- and CD42+ MK (Figure 4D, group 5), and 17.7% of circRNA followed mRNA expression changes (groups 1 and 4). Conversely, in more than half of the circRNAs (65.1%), circRNA levels increased but the mRNA decreased (Figure 4D; group 2; 14.5%) or remained unchanged (Figure DC; group 3; 50.5%). We found no circRNA with decreased expression levels and increased mRNA levels (Figure 4D, group 6). In conclusion, the expression pattern of circRNA expression only partially follows the mRNA expression, and more than half of the circRNAs change their expression pattern independently of mRNA expression during erythroid and megakaryoid differentiation.

### The majority of circRNAs expressed in differentiated cells correlates with that of mRNA expression in progenitors

Previous studies reported that circRNAs accumulate in neuronal and muscle cells upon differentiation (26, 53, 54). To test whether this accumulation is also seen during blood cell differentiation, we first compared the number of circRNAs identified in *in vitro* differentiated erythroblasts (Figure 2D) with that of our previously published data on mature blood-derived red blood cells (RBC; (4)). The overlap of circRNA expression of *in vitro* differentiated RBC with mature RBCs was only 11.5% (676 out of 5,878 high confidence circRNAs; Figure 4E, left panel). The overlap between circRNA expression in platelets (4) and MK (Figure S2F) was higher with 17.8% (1907 out of 10,729 high confidence circRNA; Figure 4E, right panel). This limited overlap could stem from the described uptake of transcripts by platelets from exogenous sources (55), and could possibly include circRNAs. Alternatively, the differential circRNA expression could derive from transcription detected in progenitor cells. To address the latter possibility, we compared the mRNAs expressed by differentiating erythrocytes and megakaryocytes with that of circRNAs expressed RBCs and platelets, respectively. To our surprise, the mRNA-expressing genes of *in vitro* differentiating cells and circRNA-expressing genes of mature cells almost completely overlapped (95.82% and 96.45% for RBC and platelets, respectively, Figure 4E). Thus, the circRNAs detected in enucleated platelets or RBCs correlate better with the mRNA expression than with that of circRNA expression in their respective progenitors. This finding also suggests that circRNA uptake from the environment may not be the primary source of circRNA expression discrepancy between progenitors and mature cells.

### CircRNA expression does not correlate with changes in translation efficiency of the mRNA counterpart

A recent study showed that the YAP circRNA prevents the translation of its mRNA counterpart in a sequence-specific manner (22). Specifically, over-expression of YAP circRNA prevented the translation initiation complex from engaging with the YAP mRNA (22). This finding prompted us to determine the transcriptome-wide correlation of endogenous circRNA expression with the translation efficiency of their mRNA counterpart. We used normalized expression data from the matched RNA-seq and ribo-seq analysis of CD42- and CD42+ MKs (Table S4). The ribosome footprint (RFP) reads fulfilled all quality controls upon mapping the reads onto the genome, i.e. the read size distribution, periodicity and reads distribution after P-site correction (Figure S3). We determined whether mRNA expression related to changes in RFP abundance by computing the LFC in mRNA abundance and the LFC in RFP abundance for each gene, between CD42- and CD42+ MK (Figure 5A, Table S4). Most changes in mRNA expression correlated well with changes in RFP counts (Pearson’s correlation coefficient: 0.854). In addition, genes that expressed circRNAs (red dot) or not (gray dots) showed very similar patterns (Pearson’s correlation coefficient: 0.768). This finding suggests that the ribosomal occupancy of mRNAs was not influenced by the co-expression of circRNA variants. Similarly, when we calculated the translation efficiency per gene *(normalized RFP counts (TPM) / normalized mRNA-seq (TPM))* (Figure 5B), no differences in mRNA translation were observed between genes that do (red dots) or do not (grey dots) express circRNAs. Notably, 1942 mRNA-expressing genes (5.49%) altered their translation efficiency with LFC>2 between CD42- and CD42+ MKs, while only 4 out of 1544 mRNA-expressing genes that also expressed circRNA (0.25%) displayed the same behavior (Figure 5B, Table S4).

**Figure 5:**
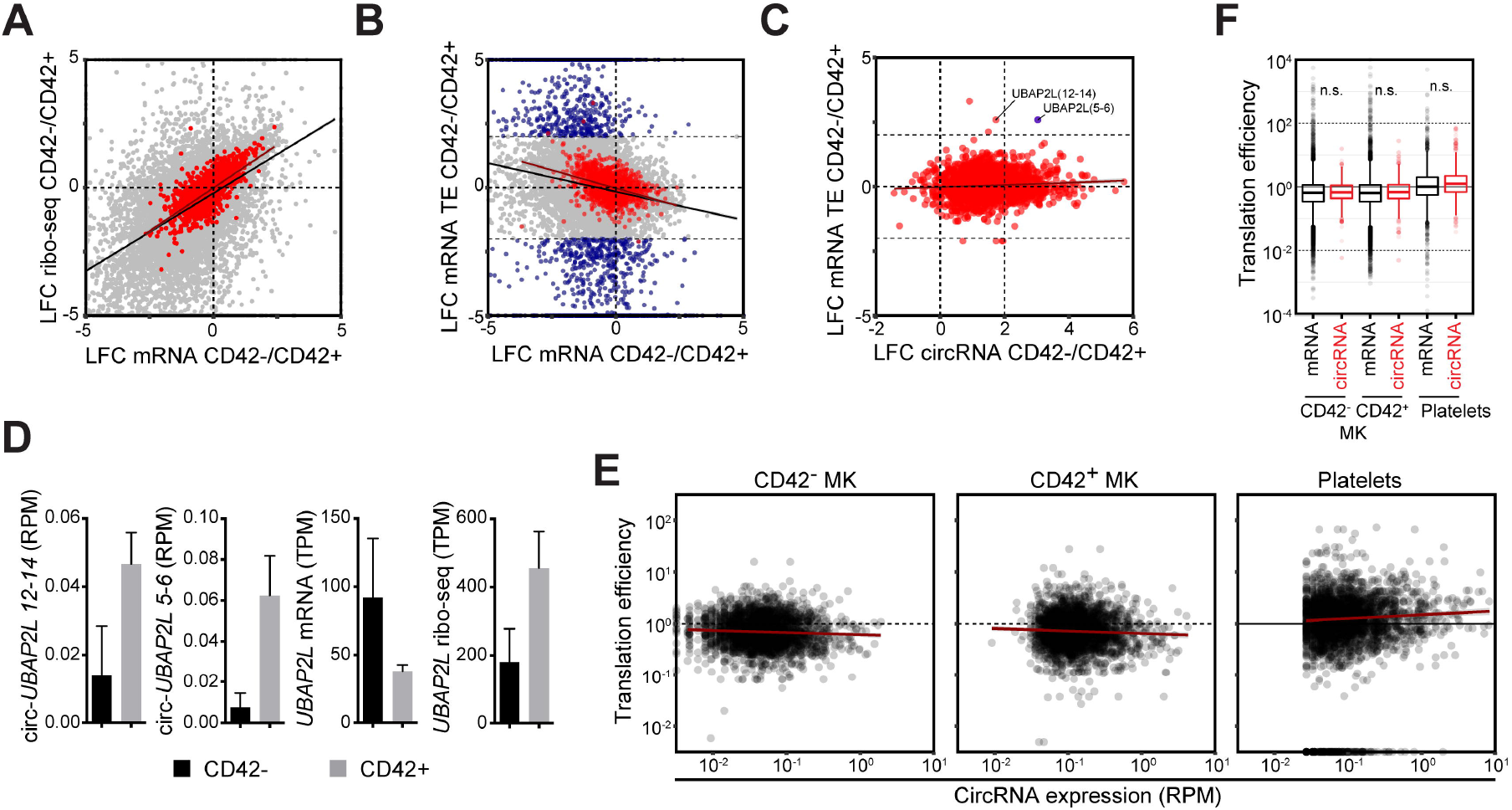
CircRNA-mediated regulation of mRNA translation is limited in megakaryocytes. (**A**) Log2 Fold Change (LFC) in ribosomal occupancy (ribo-seq) and mRNA expression between CD42- and CD42+ megakaryocytes. CircRNA-expressing genes are plotted in red. (**B**) LFC in translation efficiency (TE;see methods) and mRNA expression between CD42- and CD42+ megakaryocytes. (**C**) LFC of translation efficiency and circRNA expression between CD42- and CD42+ megakaryocytes. Genes are plotted in blue in B-C when LFC in translation efficiency was more than four-fold (LFC>2). (**D**) CircRNA, mRNA and ribo-seq expression of *UBAP2L*. (**E-F**) Relation between circRNA expression and translation efficiency (**E**) and translation efficiency for circRNA- and mRNA-only expressing genes (**F**) in CD42-, CD42+ and platelets. (A-C) The linear regression of all genes is represented inblack, and this of circRNA expressing gene in dark-red. RPM: reads per million mapped (linear) reads, TPM: transcripts per kilobase per million.

We next determined whether changes in translation efficiency of mRNAs upon MK maturation related to the expression of circRNA from the same gene. Overall, we observed lower changes in translation efficiency in circRNA-expressing genes than circRNA-less mRNA-genes (span of LFC in translation efficiency = 5.42 and 29.55, respectively; Figure 5B). Only 1 circRNA (2.67%), circ-UBAP2L, altered its expression (LFC>2) in CD42+ compared to CD42- MKs and showed an alteration in translation efficiency (LFC>2) of the mRNA counterpart (Figure 5C; blue dot). Closer examination of the mRNA, RFP and circRNA levels of UBAP2L gene indeed suggested a correlation of changes in translation efficiency with circRNA expression (Figure 5D).

To further examine whether the lack of abundant circRNA association with mRNA translation efficiency was specific to MKs or more broadly applicable, we examined the correlation of translation efficiency with circRNA expression throughout platelet maturation (Figure 5E). We found that the translation efficiency in CD42-, CD42+ and platelets remained uncorrelated to circRNA expression. Furthermore, the overall mRNA translation efficiency remained unchanged, whether the genes expressed circRNAs or not (Figure 5F). Notably, this lack of association between circRNA and changes in translation efficiency of the mRNA counterpart was also observed in K562 and HeLa-S3 human cell lines (Figure S4). Combined, these findings indicate very limited effects of circRNAs on the translation efficiency of their mRNA counterpart.

### CircRNAs contain putative open reading frames

Most circRNAs (~80%) are reported to fall in the coding region and/or contain the canonical translation start site (5, 19). We therefore sought to define the coding potential of megakaryocytic circRNAs. We used ORF-finder (NCBI) to predict open reading frames (ORFs) in high confidence circRNAs. Because algorithms such as ORF-finder cannot mimic back-splicing in silico, we juxtaposed the circRNA sequence three times (termed 3xCircRNA; Figure S5A). This 3xCircRNA sequence allows the investigation of ORFs spanning over the back-splicing junction that could gain stop codon as a result of frameshifts at the circRNA back-splicing junction. To identify ORFs that are common to both mRNAs and circRNAs, we also used the high confidence circRNA sequence opened-up at the back-spliced junction (1xLinRNA; Figure S5A). Open reading frames with a minimal length of 25aa and a canonical AUG start codon in the 1xLinRNA and the 3xCircRNA sequences were included in the analysis.

The majority of 1xLinRNA sequences (2,371 out of the 2,531) in MKs contained putative ORFs (n= 8,519 unique ORFs; mean ORF length: 77.3 aa; median: 46aa). Similarly, 2,488 out of 2,531 3xCircRNA contained at least one predicted ORF (n= 12,567 unique ORFs; mean ORF length: 115.9aa; median: 55aa; Figure 6A). To isolate circRNA-specific ORFs, we subtracted the ORFs found in 1xLinRNA from those found in 3xCircRNA sequences. Whereas 7,559 unique ORFs were shared by both sequences (mean ORF length: 66.99aa; median: 42aa), 5,008 unique ORFs were specific for 2,428 circRNAs (mean ORF length: 189.6aa; median: 109aa; Figure 6B). CircRNA-specific ORFs were longer than those shared by 1xLinRNA and 3xCircRNA (p-adjusted= 1.24e-12; Figure S2G). Thus, 95.93% of circRNAs in MKs have specific putative open reading frames. CircRNA-specific putative ORFs were also detected in platelets, with 19,995 ORFs (6,904 out of 10,729 circRNAs, 64.3% of circRNA), and in differentiating erythroblasts, with 1,864 ORFs (639 out of 950 circRNAs, 67.3% of circRNA; Figure S5B-C).

**Figure 6:**
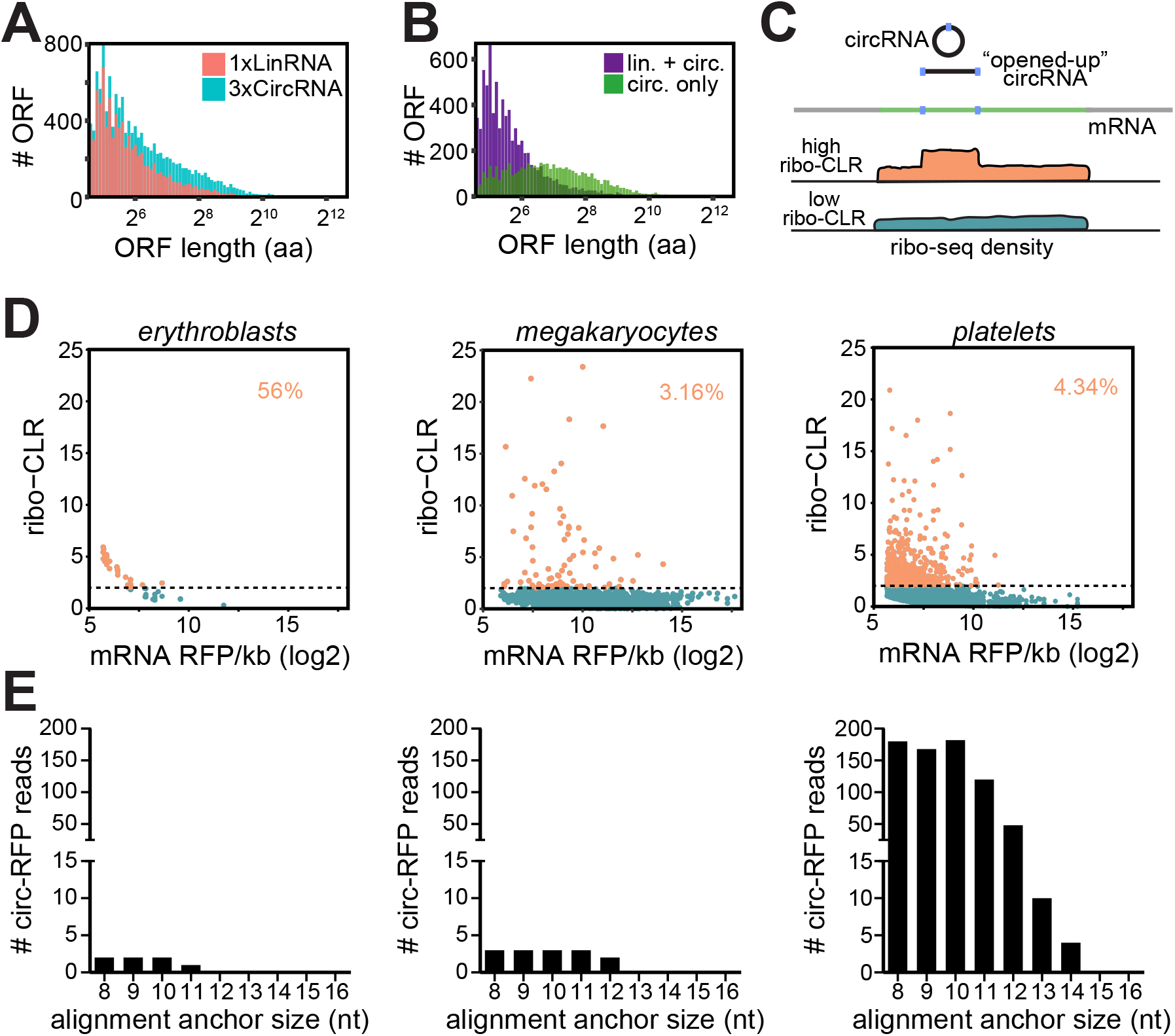
Putative ORF detection in circRNAs, with limited evidence for translation. (**A**) Open reading frames (ORFs) were computed (see methods) for linearized high confidence circRNA sequences (1xLinRNA), and for tripled juxtaposed linearized sequence of high confidence circRNA in megakaryocytes to include circularized junction areas (3xCircRNA, see Figure S4A). Bar graph depicts the length and frequency of all identified ORFs found in 1xLinRNA (pink) or 3xCircRNA (blue). (**B**) Bar graph depicting the length and frequency of the ORFs found in both 1xLinRNA and 3xCircRNA sequences (purple) or specifically in circRNA (green). (**C-D**) Ribosome footprinting sequencing (Ribo-seq) data were used to determine the ribosome density per kilobase on mRNA sequence and circRNA sequence. The circular-over-linear ribosome density ratio (ribo-CLR) was calculated. (**C**) Schematic representation of circRNA (back-spliced junction in blue) and mRNA sequence, and ribo-seq density resulting in high or low ribo-CLR. (**D**) Calculated ribo-CLR for erythroblasts (left panel), megakaryocytes (middle panel), and platelets (right panel) and plotted against the mRNA ribosome footprint (RFP) density (RFP per kilobase). (**E**) Ribo-seq data of erythroblasts (left panel; data from (42)), CD42- or CD42+ megakaryocytes (middle panel), and platelets (middle panel; data from (42)) were screened for RFP reads on the circRNA junction (circ-RFP) using CE2. The minimum chimeric alignment anchor size for alignment (8 to 16 nt) is indicated. Circ-RFP that were not found expressed in low confidence circRNA (based on RNA-seq detection) were excluded.

### A subset of circRNAs displays a high ribosome density

To determine whether the circRNA-containing putative ORFs show any evidence of translation into proteins, we re-analyzed previously published RFP sequencing data from erythroblasts (42). We determined the ribosome density on the coding region of mRNA sequence, and on the circRNA sequence (Figure 6C). We mapped the RFP read on the whole mRNA coding sequence and on the ‘high confidence’ circRNA sequence that was opened-up at the back-spliced junction. We then calculated the RFP density per kilobase on the mRNA and the circRNA sequences. To measure differences in RPF reads between circRNA and mRNA, we calculated the ratio of circRNA-over-mRNA RFP reads density (ribo-CLR). Of note, a ribo-CLR > 1 would be indicative of a higher RFP density over the circRNA sequence than on the mRNA sequence. Erythroblasts contained only 57 circRNA-mRNA ribo-seq pairs that had reliable expression levels (>50 RFP per kilobase; Figure 6D, left panel; Table S4). Out of these 57 circRNAs, 32 (56% of pairs) had a ribo-CLR >2. It is thus conceivable that these 32 circRNA are translated in erythroblasts.

In MK ribo-seq data, the vast majority (96.84%) of the circRNA-expressing genes had a ribo-CLR below 2, which indicates similar mRNA and circRNA RFP read densities (Figure 6D, middle panel; Table S4). Indeed, of the 2,531 circRNAs, only 80 (3.16%) showed a higher RFP density than their corresponding mRNA (Table S4). Lastly, we re-analyzed published ribo-seq data of platelets (42). 465 circRNAs showed a ribo-CLR>2 (Figure 6D, right panel; Table S4). Nevertheless, with 4.34% of all 10,729 circRNAs, the overall percentage of circRNAs that could potentially contribute to the platelet proteome resembled that of MK. Thus, a small proportion of circRNAs in erythrocytes, megakaryocytes and platelets showed a higher ribosome density than their mRNA counterpart.

### CircRNAs in platelets show hundreds of ribosome footprints on the back-spliced junction

Ribosome occupancy should also be found at the back-spliced junction of the circRNA, if this region contributes to the putative circRNA-specific coding region. We therefore re-screened all RFP reads from erythrocytes, megakaryocytes, and platelets and with an adaptation of the ribo-seq reads alignment to allow for chimeric reads detection and subsequent back-spliced junction detection. Overlap of the chimeric reads to the genome was set with a minimum anchor point of 8nt to 16nt on either side of the read. We also allowed 1 mismatch. Of the 114 million reads in erythroblasts, we found only 2 RFP reads for anchors of 8, 9, or 10nt (Figure 6E, left panel; Table S4). The 2 RFPs corresponded to 2 circRNAs *circ-ARHGEF12* and *circ-SPECC1*. As the median ribo-seq read density in erythroblasts reached 1.984 ×10^−3^ reads per nt, one would already expect by chance about 53 reads on the 950 ‘high confidence’ circRNA (0.055 read/circRNA junction).

We also screened the 757 million ribo-seq reads in megakaryocytes for ribosome footprints at the back-spliced junction. As the median ribo-seq read density for all translated genes in megakaryocytes was 0.105 reads per nt, one would expect by chance about 7,428 reads on the 2,531 ‘high confidence’ circRNA if they were translated (2.93 reads/circRNA junction). However, only 3 RFPs reads matching circRNAs in megakaryocytes were detected (Figure 6E, middle panel; Table S4). These 3 RFPs reads corresponded to 2 circRNAs; i.e. *circ-PRDM2 and circ-YEATS2*.

Of the ~178million RFP reads in platelets, one would expect by chance about 3378 RFPs on the 10,729 ‘high confidence’ circRNA of platelets (median RFP density on mRNA 0.011read/nucleotide, 0.314 read/circRNA junction). Yet, we only detected 180 RFPs, 168 RFPs and 182 RFPs, over the circRNA back-splice junction in platelets with an anchor size of 8, 9, and 10nt, respectively (Figure 6E, right panel; Table S4). The circRNA-RFPs corresponded to 56 different circRNAs. Of note, 13 of these 56 circRNAs (*HIPK3, TMEM135, CORO1C, DNAJC6, GSAP, ASH2L, FAM120A, MCU, FARSA, WDR78, ZC3H6, NCOA2, APOOL*) also displayed a ribo-CLR>2 (Figure S5D), indicating ribosome reads on both the back-splice junction and the full circRNA sequence. Overall, the RFP analysis showed some but limited evidence of translation, in particular in MK, and erythroblasts. Platelets provide the best indication of putative circRNA translation, with - based on the sequencing depth of the riboseq data sets- ~58 times more circRNA-specific ribo-seq reads than erythroblast and ~258 times more than megakaryocytes.

### Mass spectrometry fails to identify circRNA-specific peptides in platelets

Because the RFP analysis in platelets showed the highest probability of translation, we further searched for evidence of translation in this cell type. We generated a reference peptide library from the 3 translation frames over the 508 circRNA back-spliced junction of the high-confidence circRNAs with ribo-CLR above 2 or detected circ-RFP. Translation frame(s) that contained a STOP codon before the back-spliced junction were excluded from this reference peptide library. This left 1566 unique putative circRNA peptides spanning the back-spliced junction with a minimum of 5 amino acids overhang. Of these 1566 putative circRNA peptides, 12 (0.77%) did not contain lysine nor arginine, 97 (6.2%) and 123 (7.9%) contained only arginine, or lysine, respectively. Thereby the majority (99.2%) of putative circRNA peptides are cleavable by trypsin treatment and should be detectable by mass spectrometry. To also identify circRNA-specific peptides outside the circRNA-junction, we included the full-length circRNA-specific putative ORFs of platelets in the peptide library we identified in Figure S4B. Lastly, we also included the reference proteome (UniProt) as decoy, to prevent possible ‘background’ hits to match known canonical protein sequences. Published MS data of platelets (44) were used to search for circRNA-specific peptides. Of the 192,743 identified peptides, not one peptide matched a circRNA junction (Figure S5D). Thus, even though ribo-seq data suggest some possible translation from circRNAs in platelets, the detection of peptides from back-spliced circRNA junctions, or from circRNA-specific ORFs fall- if present at all- below the detection limit of data-dependent Mass Spectrometry analysis.

## DISCUSSION

In this study, we present a comprehensive analysis of circRNAs in terminal differentiation of erythroblasts and platelets. CircRNA expression is in particular increased at enucleation in erythroid differentiation. This could reflect an accumulation of transcript degradation left-overs upon enucleation, as previously suggested (30). While this is an attractive hypothesis, not all circRNA follow this overall trend. In fact, more than 50% of circRNAs were regulated independently from changes in linear mRNA expression. This was true when we compared differences in expression levels prior to and post enucleation. We also observed a clear increase in number and expression levels of specific circRNAs at enucleation, which were not detected in progenitors. This increase of circRNA number and amount is unlikely to stem from a technical reason, as all samples had sufficient sequencing depth (~30 million reads) to allow for a reliable detection of lowly expressed circRNA. Thus, the majority of measured circRNAs in terminally differentiated erythrocytes cannot be explained by mere accumulation from progenitors alone. This was also true for circRNA expression in MK compared to platelets.

Another possible explanation for the increase in circRNA during erythroid maturation could be the loss of the circRNA degradation machinery. Even though it is not fully understood how circRNAs are degraded, the RNAse-L, G3BP1 and G3BP2 have been implicated in this process (56, 57). Notably, G3BP1 & 2 are down regulated during erythroid maturation, and RNAse-L is absent in mature erythrocytes (58). It is therefore tempting to speculate that the loss of the circRNA degradation machinery in mature erythrocytes contributes to the increased circRNA levels.

Intriguingly, the circRNAs in differentiated RBCs and platelets strongly overlapped with the mRNAs that were expressed in their progenitors. It has been proposed that splicing occurs in the cytoplasm of platelets and MK (59, 60). It is therefore tempting to speculate that back-splicing and formation of circRNA could also occur in the cytoplasm. In line with this, several splicing factors were detected in platelets by MS analysis (44, 61)), including DHX9. Therefore, at the period of enucleation of erythrocytes and during platelet formation, mRNA could be circularized, and thus contribute to increased numbers of circRNAs during enucleation and therefore explain the substantial overlap with the mRNAs in progenitors. This hypothesis, however, requires experimental confirmation. In addition, MK can sort specific mRNA into platelets (62). Whether circRNA can also be sorted and whether this sorting could explain some of the discrepancy between MK and platelets circRNA content, is yet to be uncovered.

About 4% of circRNAs in platelets and erythrocytes did not overlap with the mRNA levels in MK and differentiating erythroblasts. These circRNAs could be acquired from other cell types in the circulation, as observed for linear RNA in the context of tumor ‘educated’ platelets (55), or endogenous vascular-derived RNA (63). A study in platelets from healthy and diseased individuals could shed light on this. If true, circRNAs taken up by platelets or erythrocytes could in fact serve as novel biomarkers, which could be in particular valuable for diagnostics due to their high stability.

The function of circRNAs still remains elusive. CircRNA could serve to reinforce transcription, and lead to differential exon usage as previously proposed (64). Differential exon usage was also recently reported to be supported by pre-mRNA exonic sequences (65). Whether this is indeed occurring in primary blood cells is yet to be determined.

CircRNAs are also reported to regulate mRNA translation in a sequence-specific manner (22–24). We investigated here the possible effect of circRNA on its mRNA counterpart. We only identified 1 circRNA in MKs that show limited indications for translation regulation. In addition, we found no correlation between translation efficiency of mRNA and the expression level of their circular counterpart, whether in MK, platelets, or human cell lines. Whether our findings in MKs are also applicable to enucleated erythrocytes is not known. CircRNA-mediated regulation of mRNA translation is an attractive hypothesis in erythrocytes, because the majority of translation is shut down in mature cells (42), and circRNAs could support translation shutdown. In addition, it is also conceivable that circRNA affect translation of other mRNA (23, 24).

CircRNAs were also shown to be translated (25–28, 66), in particular from exogenously expressed circRNAs. This finding suggests that the highly stable circRNA transcripts could help maintain the proteome in long-lived red blood cells and platelets. However, we found very limited indications that translation from endogenous circRNAs occurs. Similar to a recent study (19), we detected only few circRNA-specific RFP. The low number of detected circRNA-specific RFP reads could not be explained by low sequencing depth, as the ribo-seq datasets entailed 114, 757, 178 million reads in erythroblasts, MKs, and platelets, respectively. Yet, only platelets had 180 RFP reads on the circRNA back-spliced junction. Nevertheless, whether the increase of circRNA-specific RFP stems from active translation or from the loss of regulators of ribosome homeostasis such as PELO by erythrocytes and platelets (42) is yet to be determined.

Deep MS analysis on platelets did not identify peptides from circRNA back-spliced junctions. Yet, circRNA-derived proteins may be generated on low levels and/or result in production that is below the detection limit of MS. This finding is corroborated by recent ribosome footprint analysis that found very limited translation of >300.000 circRNAs (0.73% of circRNAs), and closely matches our findings (0.52%; 56 out of 10729 circRNAs in platelets). Indeed, as circRNA peptides are only found in minute abundance, their detection was shown to be indistinguishable from noise in MS data (67). Nonetheless, immunoprecipitation enrichment of specific targets could help to overcome this detection limit for specific circRNA (27, 28). mRNA translation also may mask the circRNA translation and its contribution to the proteome, when the circRNA-derived peptides are identical to the mRNA-derived peptides. Nevertheless, because only 117 out of the 465 circRNAs (25.16%) with high ribo-CLR show protein expression in MS, the relative contribution of circRNA-derived translation is arguably limited in light of the mRNA-derived translatome. Compiled, our study thus only supports a limited possible contribution of circRNA-translation to the proteome in platelets.

In conclusion, we here provide the landscape of circRNA expression during in erythroid differentiation and megakaryocyte maturation. This study can serve as blueprint for future studies in these cell types. Our results also challenge the current views on the ubiquitous functions proposed for circRNAs in the biological context of terminal hematopoiesis.

## DATA AVAILABILITY

Ribosome footprinting and RNA-seq dataset of megakaryocytes were deposited on NCBI’s Gene Expression Omnibus (GEO) with accession number GSE159579. Scripts used in this study are accessible on GitHub (https://github.com/BenNicolet/CircRNA_in_MK_and_Erys).

## ACKNOWLEDGMENTS

We also thank S. Heshusius for helpful discussions, and M. Hansen, P.J. van Alphen, and M. van den Biggelaar for critical reading of the manuscript.

## AUTHOR CONTRIBUTIONS

B.P.N. analyzed data, B.P.N., S.J, and E.H. performed experiments. M.C.W., W.H.O, E.v.d.A, and M.v.L supervised, B.P.N. and M.C.W wrote the manuscript.

