## Supplementary figure legend for "Circular RNAs exhibit limited evidence for translation, or translation regulation of the mRNA-counterpart in terminal hematopoiesis"

**Figure S1: expression of high confidence circRNA and their relation to mRNA in erythropoiesis.**

(**A**) Principal component analysis (PCA) of mRNA expression of erythroid differentiation used in Figure 1 and megakaryoid differentiation used in Figure 3, also including relevant cell populations (see Table S1). (**B**) Gene ontology analysis of circRNA-expressing genes found in differentiating erythroid cells (high confidence circRNA; n=950), for biological processes (left panel) and molecular function (right panel). Bars in red indicate a negative enrichment. (**C**) Heatmap of high confidence circRNA expression during erythropoiesis (RPM; n=950). Color scale represents Z-score. (**D**) *Circ-FIRRE* expression level at indicated days of erythroid differentiation. (**E**) Circular-over-linear ratio (CLR) was calculated for all high confidence circRNA. CLR was plotted against the circRNA expression (RPM: Reads per million mapped reads) at the indicated day of differentiation.

**Figure S2: Characteristics of circRNA expression in megakaryocytes**

(**A**) Number of genes encoding mRNA, ncRNA and circRNA detected during megakaryocyte maturation and compared to platelets (cut-off for mRNA and ncRNA >0.1 TPM; low confidence circRNA). (**B**-**C**) Characterization of low confidence circRNA putative exon usage (**B**) and putative spliced length (**C**), using splicing annotations from the linear isoforms. (**D-E**). Characterization of start (**D**) and end (**E**) exon usage of ‘low confidence’ circRNAs, based on the circRNA junction positions and canonical mRNA splicing annotations. Percentage indicated the fraction of circRNA in the bar. (**F**) CircRNA detection of high confidence circRNAs. (**G**) Gene ontology analysis of circRNA-expressing genes found in differentiating megakaryocytes (high confidence circRNA; n=2531), for biological processes (left panel) and molecular function (right panel). Bars in red indicate a negative enrichment. (**H**) Open reading frames (ORFs) were computed (see method) for linear ‘opened-up’ sequence of circRNAs (1xLinRNA), and for tripled juxtaposed sequence of high confidence circRNA, to mimic a circular sequence (3xCircRNA; see Figure S4A). Bar graph showing the length of 1xLinRNA (LinRNA) or circRNA-specific (CircRNA) ORFs in megakaryocytes. Group difference in (G) was assessed with a two-tailed t-test followed by B-H p-value correction. ****p<0.0001.

**Figure S3: Quality control of ribo-seq data in megakaryocytes**

Ribo-seQC (Calviello et al., 2019) was used to assess the quality of ribo-seq reads of megakaryocytes. (**A**) Diagram depicting the mapped read length distribution of all 6 megakaryocyte ribo-seq samples within the coding sequence (CDS), 5; and 3’ untranslated regions (UTR), non-conding RNA (ncRNA) isoforms (nc Isoforms), ncRNAs, introns and intergenic sequences. (**B, C**) P-site position and periodicity after P-site position correction (**B**; 12nt), and (**C**) P-site profile across all read lengths. (B-C) are of CD42- megakaryocytes (replicate 1) and are representative for all 6 samples.

**Figure S4: circRNA-specific ORFs are abundant in erythroid cells and platelets**

(**A-B**) mRNA translation efficiency was calculated for (**A**) K562 and (**B**) HeLa-S3 cell lines, from previously published data (see methods), and plotted against mRNA expression (left panels) or circRNA expression (right panels). Red dots represent circRNA-expressing genes and black dots represent mRNA-only expressing genes. Regression line for circRNA-expressing gene is shown in dark-red while this of mRNA-expressing gene is shown in black.

**Figure S5: circRNA-specific ORFs are abundant in erythroid cells and platelets**

(**A-C**) Open reading frames (ORFs) were computed for high confidence circRNA sequences that were linearized at the back-spliced junction (1xLinRNA), and for tripled juxtaposed linearized sequence of high confidence circRNA to include circularized junction areas (3xCircRNA). (**A**) Schematic representation of the circRNA, 1xLinRNA and 3xCircRNA. Start (green) and stop (red) codon position can results in LinRNA + circRNA (Lin. + circ.) ORFs (purple), or circRNA-only (circ. only) ORFs (green) when the ORFs span over the back-spliced junction (blue), which can results in back-slice induced frame-shift (orange). (**B-C**) Bar graph depicting the length and frequency of the ORFs found in both 1xLinRNA and 3xCircRNA sequences (purple) or specifically in 3xCircRNA (green) in (**B**) differentiating erythroblasts and (**C**) platelets. (**D**) Venn diagram showing the overlap in platelets of circRNAs with ribo-CLR >2, circRNAs with circ-RFP and circRNA-specific peptide detected by mass spectrometry (no peptide detected; MS; data from (Van Oorschot et al., 2019)).
