## Supplementary figures and images for "Circular RNAs exhibit limited evidence for translation, or translation regulation of the mRNA-counterpart in terminal hematopoiesis"

### Figure S1

#### Nicolet et al. Figure S1

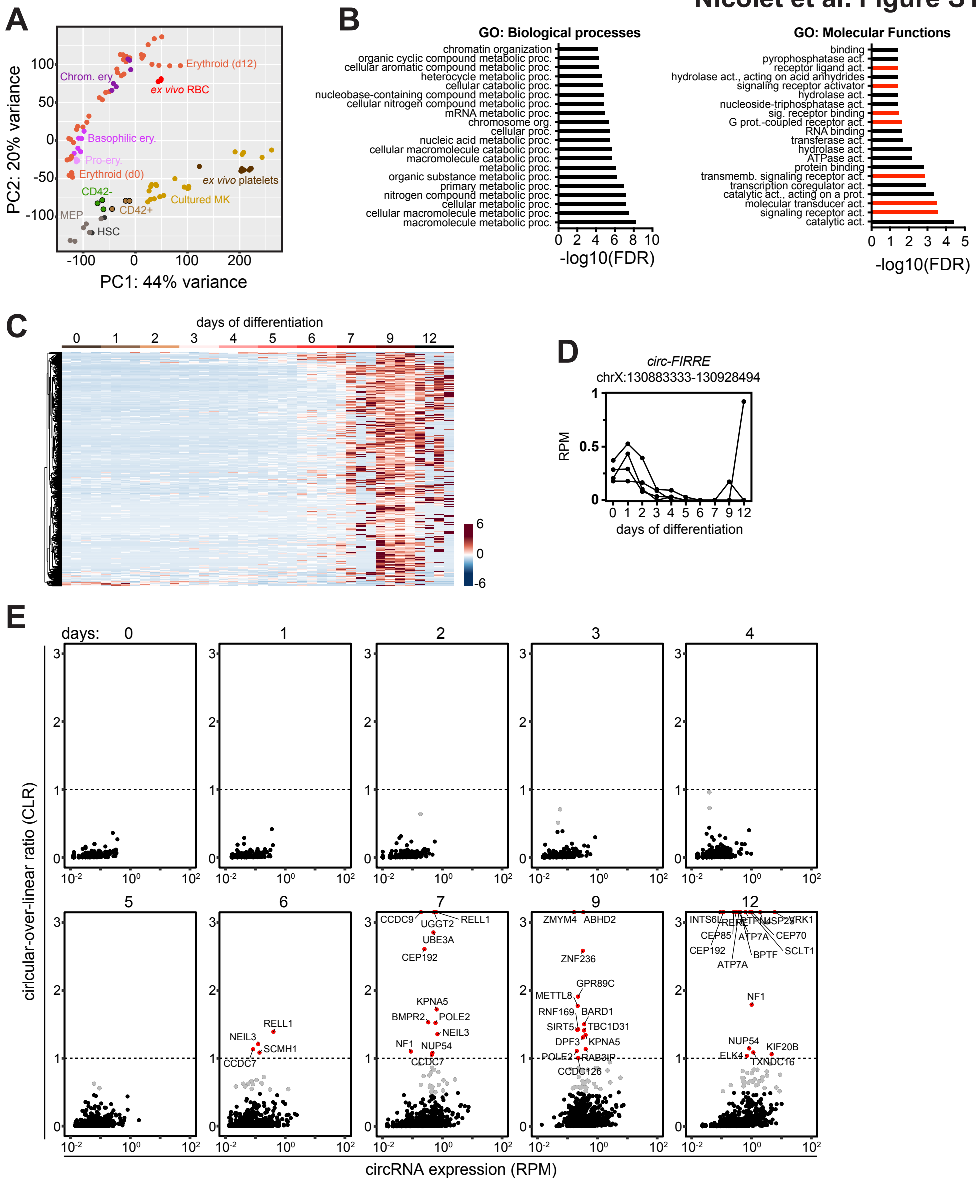

### Figure S2

#### Nicolet et al. Figure S2

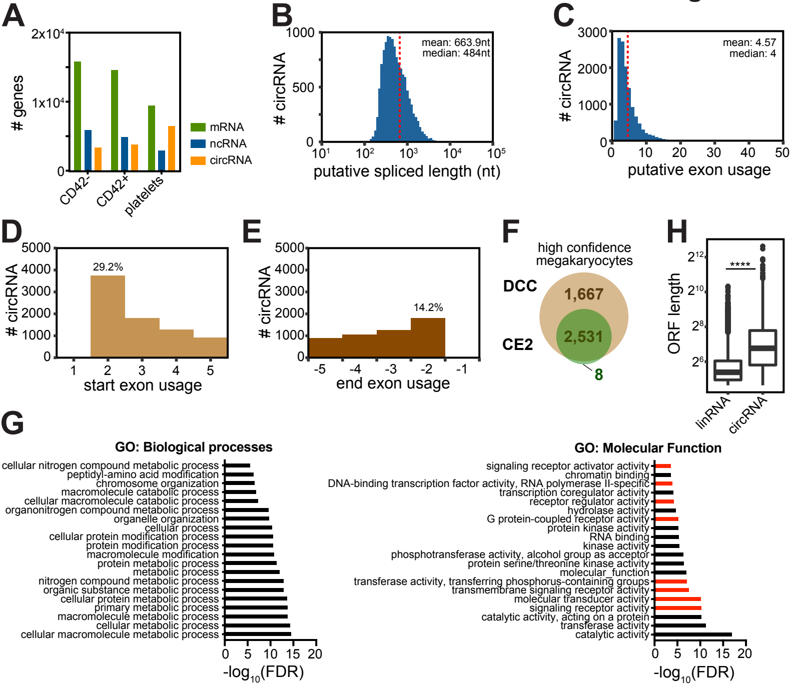

### Figure S3

#### Nicolet et al. Figure S3

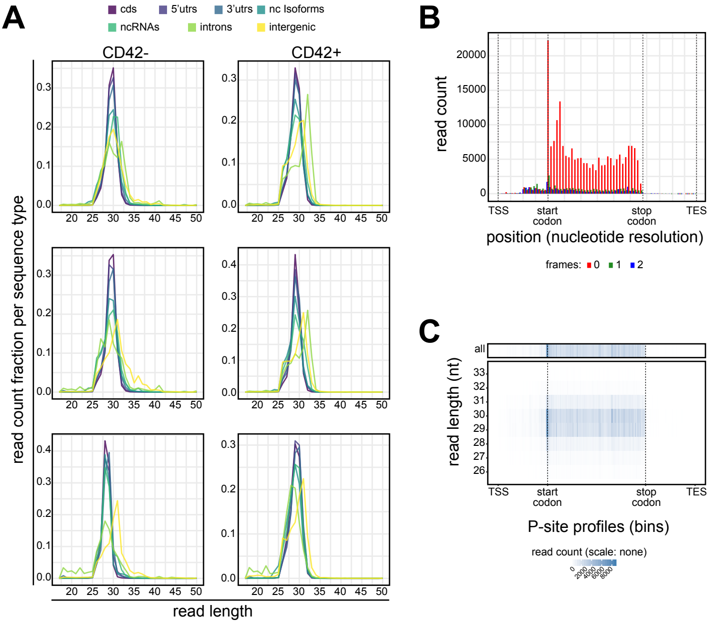

### Figure S4

#### Nicolet et al. Figure S4

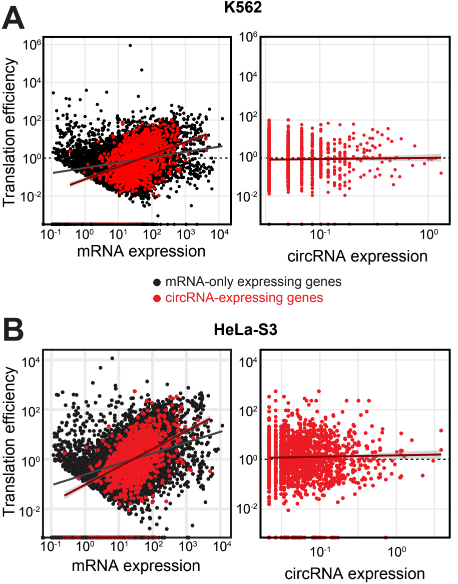

### Figure S5

#### Nicolet et al. Figure S5

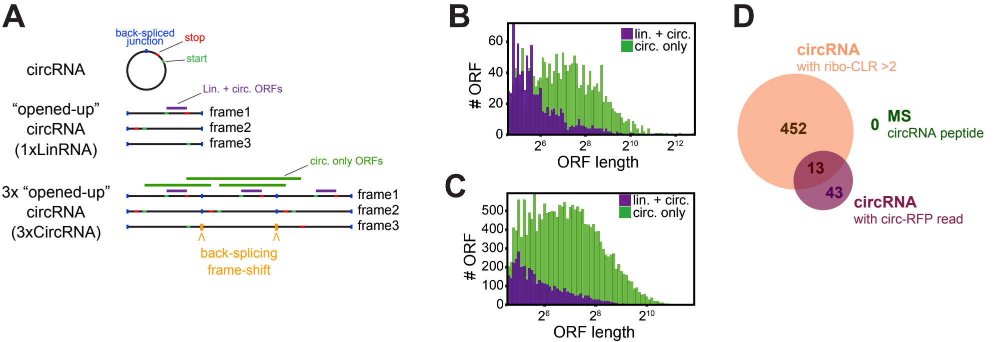
